# Influence of trunk posture on spinal loading and paraspinal muscle forces in adolescent idiopathic scoliosis: a subject-specific musculoskeletal modelling study

**DOI:** 10.64898/2026.08.28.747718

**Authors:** Rounak Bhattacharya, Bhavuk Garg, Rajesh Malhotra, Rajdeep Ghosh, Anoop Chawla, Kaushik Mukherjee

**Affiliations:** School of Interdisciplinary Research, Indian Institute of Technology Delhi, New Delhi 110 016, Delhi, India; Department of Orthopaedics, All India Institute of Medical Sciences, New Delhi 110029, Delhi, India; Currently at Department of Orthopaedics, Max Super Speciality Hospital, Saket, New Delhi 110 017, Delhi, India; Currently at Department of Orthopaedics, Indraprastha Apollo Hospital, New Delhi 110029, Delhi, India; Department of Mechanical Engineering, Indian Institute of Technology Delhi, New Delhi 110 016, Delhi, India; Currently at Department of Mechanical Engineering, National Institute of Technology Agartala, Agartala 799 046, Tripura, India

**Keywords:** Adolescent idiopathic scoliosis, Trunk posture, Musculoskeletal modelling, Spinal loading, Muscle asymmetry

## Abstract

Adolescent idiopathic scoliosis (AIS) alters spinal geometry and may influence the biomechanical response of the spine during functional postures. However, posture-dependent changes in spinal loading and paraspinal muscle forces in AIS remain poorly understood. This study investigated the effects of trunk posture on intervertebral loading and paraspinal muscle forces using a subject-specific musculoskeletal model of an adolescent with AIS. The spinal deformity was reconstructed from biplanar radiographs and incorporated into a full-body musculoskeletal model. Flexion, extension, lateral bending, and axial rotation were simulated at three incremental magnitudes, with motion distributed across the thoracolumbar spine. Intervertebral compressive and lateral forces around the curve apex and forces in the erector spinae (ES) and multifidus (MF) muscles were evaluated. Trunk flexion produced the greatest compressive loading, reaching 337 N at the curve apex and 372 N two levels below the apex at 30° flexion. Lateral bending produced pronounced direction-dependent loading: concave-side bending increased lateral forces, whereas convex-side bending increased compressive forces. Axial rotation produced similar but smaller direction-dependent changes. Paraspinal muscle forces were consistently asymmetric, with concave-side dominance of the ES and convex-side dominance of the MF. Flexion and convex-sided movements generally produced greater muscle imbalance, while increasing posture magnitude amplified spinal loading and muscle forces. These findings demonstrate that trunk posture, movement direction, and magnitude substantially influence the biomechanical environment of the scoliotic spine and should be considered when evaluating spinal mechanics in AIS.

**Highlights:**

- Spinal loading in AIS was strongly dependent on trunk posture.
- Trunk flexion produced the greatest compressive intervertebral loading.
- Concave-side bending increased lateral loading around the curve apex.
- Paraspinal muscle forces exhibited distinct concave–convex asymmetry.
- Trunk posture and spinal deformity jointly influenced AIS biomechanics.

## 1. Introduction

Adolescent idiopathic scoliosis (AIS) is a three-dimensional spinal deformity that typically develops during puberty with a higher prevalence in females (Cheng et al., 2015; Choudhry et al., 2016; Weinstein, 2019). It is characterized by lateral curvature of the spine in the coronal plane (Cobb angle >10°), accompanied by vertebral rotation and alterations in the transverse plane (Horne et al., 2014; Jada et al., 2017; Malfair et al., 2012; Weinstein, 2019). Although AIS is the most common form of scoliosis, its aetiology and mechanisms underlying curve progression remain incompletely understood (Fadzan & Bettany-Saltikov, 2017; Grauers et al., 2016; Kikanloo et al., 2019).

Trunk posture represents an important biomechanical factor influencing spinal loading during everyday functional activities. Movements such as trunk flexion, extension, lateral bending, and axial rotation are frequently performed during activities of daily living and can significantly influence paraspinal muscle recruitment patterns and intervertebral loading (Alessa & Ning, 2017; Granata & Wilson, 2001; Khoddam-Khorasani et al., 2020; Shirazi-Adl et al., 2004). In AIS, the altered three-dimensional geometry of the spine may further modify these spinal biomechanics with various trunk postures (Bassani et al., 2024; Farahpour et al., 2015; H. Wang et al., 2025; Zhang et al., 2021). However, these effects of different trunk postures on intervertebral loading and paraspinal muscle recruitment in the scoliotic spine remain poorly understood. This limited understanding is largely due to the challenges of directly quantifying spinal loading *in vivo*, as existing approaches, including intradiscal pressure measurements, intramuscular electromyography, and instrumented vertebral implants, are invasive and have limited applicability, particularly in adolescents.

Musculoskeletal (MSK) modelling provides a non-invasive approach to estimating spinal loading and muscle activation, which are difficult to quantify in vivo. Previous MSK studies in AIS have demonstrated alterations in intervertebral loading and trunk muscle recruitment associated with the scoliotic deformity (Barba et al., 2021; Bassani et al., 2021, 2024; Schmid, Burkhart, Allaire, Grindle, Bassani, et al., 2020). For example, Barba et al. (2021) reported increasing lateral shear forces and greater asymmetry in trunk muscle activation with increasing scoliosis severity during upright standing. However, biomechanical parameters derived solely from neutral upright posture have been shown to be insufficient for predicting curve progression (Bassani et al., 2021), highlighting the need to investigate spinal loading under functional postural conditions.

To the best of the authors’ knowledge, only one study (Bassani et al., 2024) has evaluated the effects of simulated trunk motion on spinal loading in AIS. However, in that study, the thoracic spine was represented as a rigid segment, with trunk motion primarily accommodated by the lumbar spine, limiting the representation of physiological intervertebral motion. In addition, the model excluded the upper limbs and their contribution to gravitational loading. These limitations highlight the need for a more anatomically and functionally representative MSK model to investigate posture-dependent spinal loading and muscle recruitment in AIS.

Therefore, this study aimed to investigate the effects of different trunk postures on spinal loading and paraspinal muscle forces in an individual with AIS using a detailed subject-specific MSK model. Flexion, extension, lateral bending, and axial rotation were simulated at increasing magnitudes by distributing intervertebral motion across the thoracolumbar spine. The resulting compressive and lateral intervertebral forces were evaluated around the curve apex, together with the magnitude and asymmetry of erector spinae (ES) and multifidus (MF) muscle forces. This approach enabled the characterization of posture-, direction-, and magnitude-dependent changes in the biomechanical response of the scoliotic spine in AIS.

## 2. Methods

### 2.1. Study participant and data collection

One female patient with AIS (age: 16 years, height: 1.59 m, weight: 55 kg, right-convex T8–T12 curve, Cobb angle: 59°, curve apex at T10) was recruited at the All India Institute of Medical Sciences (AIIMS), New Delhi, India. Spatially calibrated anterior–posterior and lateral radiographic images were acquired in the upright standing posture using the EOS imaging system™ (EOS Imaging, Paris, France). The Institutional Review Board of AIIMS, New Delhi, granted ethical approval for the study (IEC-248/04.03.2022). Written informed consent was obtained from the patient’s guardians (since the patient was a minor) for the use of anonymized radiological data for research purposes.

### 2.2. Development of subject-specific MSK model of AIS

The base musculoskeletal model was selected from a validated set of OpenSim-based full-body models developed for healthy children and adolescents (Schmid, Burkhart, Allaire, Grindle, & Anderson, 2020), corresponding to the patient’s age and sex. The thoracolumbar spine was represented using three rotational degrees of freedom at each intervertebral joint from T1/T2 to L5/S1, while translational motion was constrained. The ribs were connected to the thoracic vertebrae through single compound revolute joints, and the combined head–neck segment was connected to the T1 vertebra using a spherical joint to permit multi-axial rotational motion. The model incorporated more than 30 muscle groups, comprising a total of 552 individual muscle fascicles, enabling a detailed representation of the trunk musculature (Schmid, Burkhart, Allaire, Grindle, & Anderson, 2020).

The three-dimensional positions and orientations of the vertebrae from T1 to L5 were reconstructed from spatially calibrated biplanar radiographic images acquired using the EOS imaging system™ (EOS Imaging, Paris, France). For each vertebra, nine anatomical landmarks were manually identified, including the upper and lower vertebral corners, as well as the tip of the spinous process, in both the anterior–posterior and lateral radiographs. These landmarks were used to determine vertebral orientation in the sagittal and frontal planes by calculating the slope of the line connecting the upper and lower vertebral corners within each respective plane (Bassani et al., 2017). Intervertebral joint centers were then estimated as the centroids of the intervertebral space, defined by the region enclosed between the lower corners of the superior vertebra and the upper corners of the adjacent inferior vertebra (Bassani et al., 2017). Subsequently, the intervertebral joint spacing was defined as the distance between adjacent intervertebral joint centers.

The extracted spinal deformity was incorporated into the scaled adolescent musculoskeletal model by modifying the positions and orientations of each vertebra from T1 to L5 in both the frontal and sagittal planes, based on subject-specific radiographic measurements (Fig. 1). These adjustments altered the alignment of the rib cage and upper limb segments; therefore, their orientations were subsequently readjusted to restore their original anatomical configuration. In addition, the head-neck segment was reoriented to maintain a horizontal gaze, thereby defining a neutral upright standing posture (Schmid, Burkhart, Allaire, Grindle, Bassani, et al., 2020). To improve computational efficiency, the rib joints were locked, and the intercostal muscles were excluded from the model (Rauber et al., 2024).

**Fig. 1.**
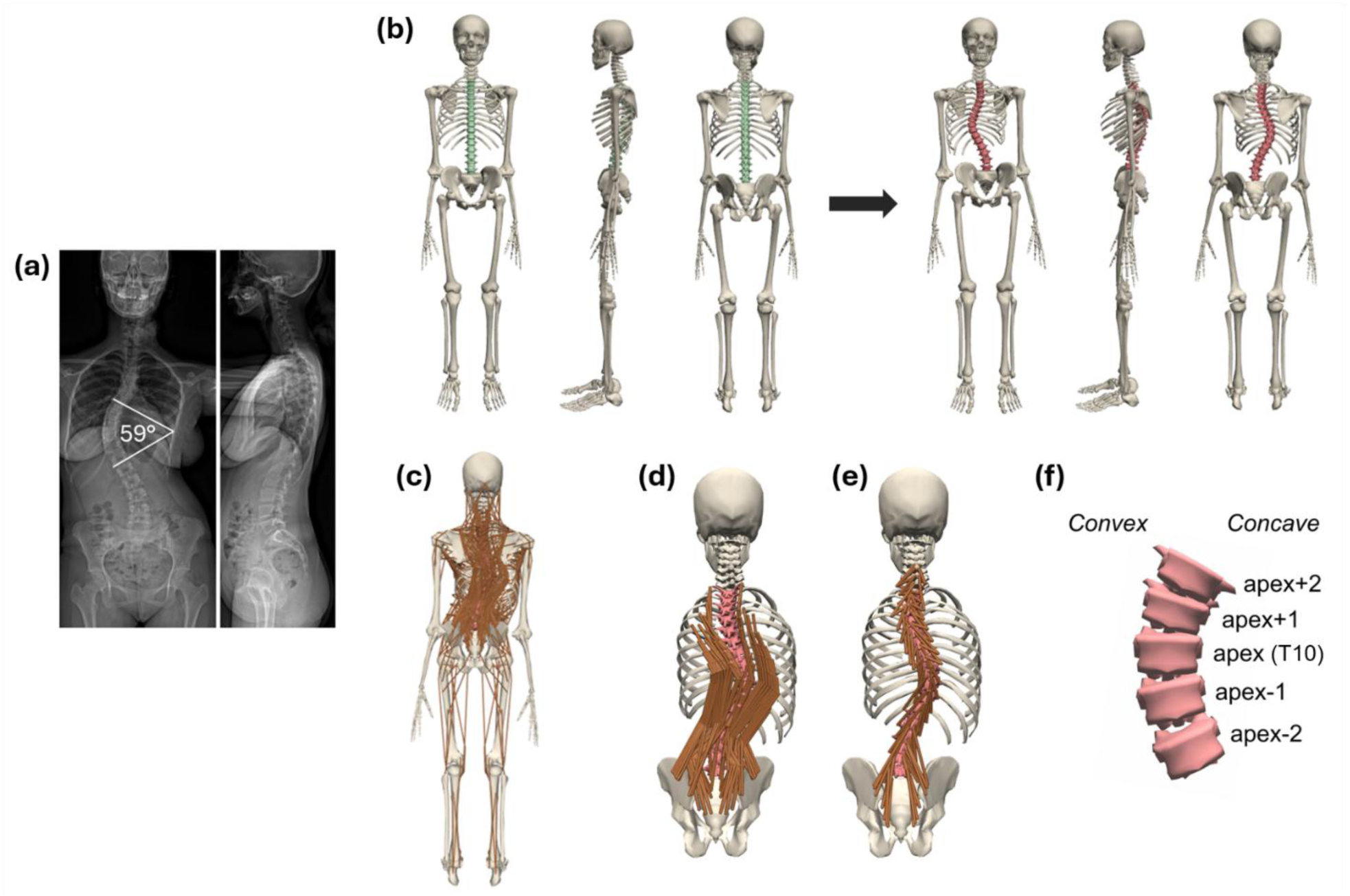
(a) EOS radiograph of the AIS patient; (b) development of the subject-specific AIS model; AIS model showing (c) all muscle groups; (d) erector spinae (ES); (e) multifidus (MF) muscle group; and (f) vertebral levels from two levels above to two levels below the curve apex (T10).

### 2.3. Simulation of trunk postures

Six standard trunk postures were simulated at three incremental degrees of rotation (Fig. 2): flexion (10°, 20°, and 30°), extension (5°, 10°, and 15°), right and left lateral bending (5°, 10°, and 15°), and right and left axial rotation (5°, 10°, and 15°). Lateral bending toward the concave (right) and convex (left) sides of the curve was defined as concave-side and convex-side bending, respectively. Similarly, axial rotation toward the concave (right) and convex (left) sides was defined as concave-side and convex-side rotation, respectively. The distribution of intervertebral motion relative to overall spinal motion was assigned based on values reported in previous studies (Fujii et al., 2007; Fujimori et al., 2012, 2014; Rozumalski et al., 2008; White & Panjabi, 1978; Wong et al., 2006), as summarized in Table 1. As intervertebral motion distribution patterns specific to scoliosis patients are not currently available, values derived from healthy individuals were adopted.

**Fig. 2.**
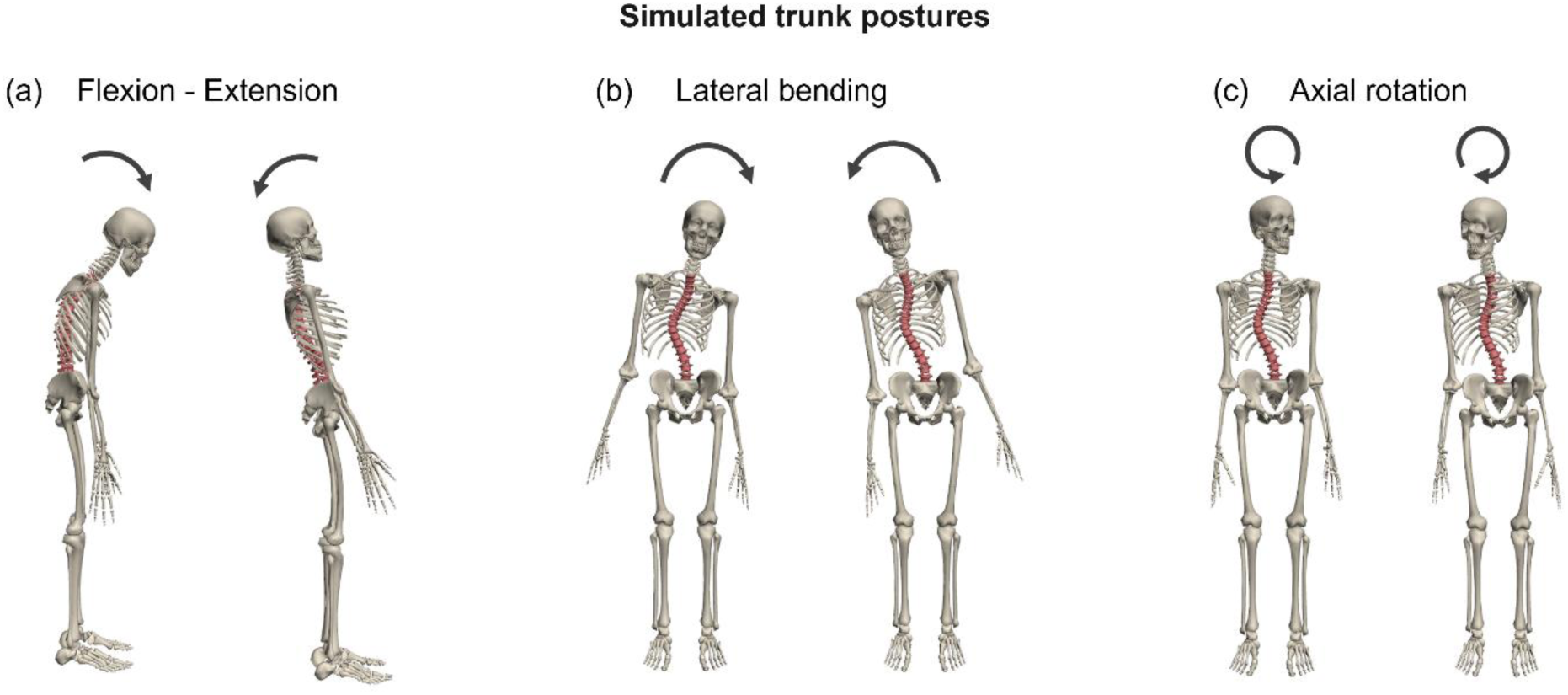
Simulated trunk postures used in the present study: a) flexion–extension, b) convex (left) and concave (right) lateral bending, and c) convex (left) and concave (right) axial rotation.

**Table 1.** Segmental contribution ratios of intervertebral motion relative to total spinal motion used to define kinematic constraints for flexion–extension (White & Panjabi, 1978; Wong et al., 2006), lateral bending (Fujimori et al., 2014; Rozumalski et al., 2008; White & Panjabi, 1978), and axial rotation (Fujii et al., 2007; Fujimori et al., 2012).

| Joint level | Flexion–extension | Lateral bending | Axial rotation |
| --- | --- | --- | --- |
| T1/T2 | 0.0265 | 0.057 | 0.046 |
| T2/T3 | 0.0265 | 0.053 | 0.062 |
| T3/T4 | 0.0265 | 0.057 | 0.054 |
| T4/T5 | 0.0265 | 0.037 | 0.062 |
| T5/T6 | 0.0265 | 0.033 | 0.070 |
| T6/T7 | 0.0330 | 0.045 | 0.074 |
| T7/T8 | 0.0395 | 0.070 | 0.089 |
| T8/T9 | 0.0395 | 0.053 | 0.097 |
| T9/T10 | 0.0395 | 0.066 | 0.105 |
| T10/T11 | 0.0590 | 0.074 | 0.101 |
| T11/T12 | 0.0790 | 0.094 | 0.050 |
| T12/L1 | 0.0790 | 0.090 | 0.019 |
| L1/L2 | 0.1500 | 0.051 | 0.029 |
| L2/L3 | 0.1235 | 0.086 | 0.031 |
| L3/L4 | 0.1055 | 0.066 | 0.038 |
| L4/L5 | 0.0735 | 0.049 | 0.038 |
| L5/S1 | 0.0340 | 0.037 | 0.036 |

An inverse dynamics-based simulation pipeline was applied to estimate muscle forces and joint reaction forces across the simulated trunk postures (Rajagopal et al., 2016; Seth et al., 2011). Muscle forces were computed using a static optimization algorithm (Erdemir et al., 2007) that minimizes the sum of squared muscle activations as the objective function. Since the influence of scoliosis on muscle physiological properties is still not well-documented, muscle force–length– velocity relationships were not incorporated into the analysis. To represent the interaction between the feet and the ground, residual actuators were applied between the calcaneus and the ground. A maximum actuator force of 10 kN was specified to provide sufficient mechanical support while minimizing the influence of residual actuators on the predicted muscle forces (K. Burkhart et al., 2020; K. A. Burkhart et al., 2018). Subsequently, joint reaction analysis was performed to calculate the intervertebral reaction forces at each spinal level.

### 2.4. Prediction of intervertebral reaction forces and trunk muscle forces

The intervertebral reaction forces were calculated at the curve apex, as well as at two vertebral levels above and below the apex. Specifically, the compressive (***F****_comp_*) and lateral (***F****_lat_*) force components were extracted and compared across the simulated trunk postures.

Erector spinae (ES) and multifidus (MF) muscle forces were evaluated on the convex and concave sides of the scoliotic curve. Total muscle force on each side was calculated by summing the forces of all muscle fascicles spanning from T1 to L5 (Fig. 1d and 1e). In addition, level-wise muscle forces were determined by summing the forces of individual muscle fascicles crossing the corresponding vertebral mid-plane (Bruno et al., 2015). Muscle force asymmetry was quantified as the difference between the muscle forces on the convex and concave sides of the curve (convex − concave). Values close to zero indicated a relatively symmetrical distribution of muscle forces between the two sides, whereas positive and negative values indicated greater muscle forces on the convex and concave sides, respectively, and thus, asymmetry in paraspinal muscle forces.

### 2.5. Model validation

Model validation was performed by comparing the predicted erector spinae (ES) muscle activation with previously reported *in vivo* measurements (Kwok et al., 2015) and MSK modelling predictions (Barba et al., 2021; Schmid, Burkhart, Allaire, Grindle, Bassani, et al., 2020) in individuals with AIS. Specifically, the convex-to-concave muscle activation ratio was used as the primary validation metric. At each vertebral level, cumulative muscle activation was calculated by summing the activations of individual muscle fascicles crossing the corresponding vertebral mid-plane. Muscle activation values ranged from 0 (no activation) to 1 (maximal activation). Activation ratios were evaluated for the thoracic and lumbar regions and at the curve apex and two levels above and below. Intervertebral reaction forces were not validated because corresponding *in vivo* measurements are currently unavailable for individuals with AIS.

## 3. Results

### 3.1. Model Validation

The predicted ES muscle activation ratios were generally consistent with previously reported in vivo measurements (Kwok et al., 2015) and MSK modelling studies (Barba et al., 2021; Schmid, Burkhart, Allaire, Grindle, Bassani, et al., 2020), as shown in Fig. 3. In the thoracic and lumbar regions, the present model predicted convex-to-concave activation ratios of 0.96 and 0.87, respectively, indicating relatively symmetrical activation with slight concave-side dominance. At and around the curve apex, the predicted ratios ranged from 0.93 to 1.03 and were comparable to those reported in previous MSK studies (Barba et al., 2021; Schmid, Burkhart, Allaire, Grindle, Bassani, et al., 2020). Overall, the predicted activation ratios were within the physiological range (Fig. 3) reported in previous in vivo (Kwok et al., 2015) and MSK modelling studies (Barba et al., 2021; Schmid, Burkhart, Allaire, Grindle, Bassani, et al., 2020).

**Fig. 3.**
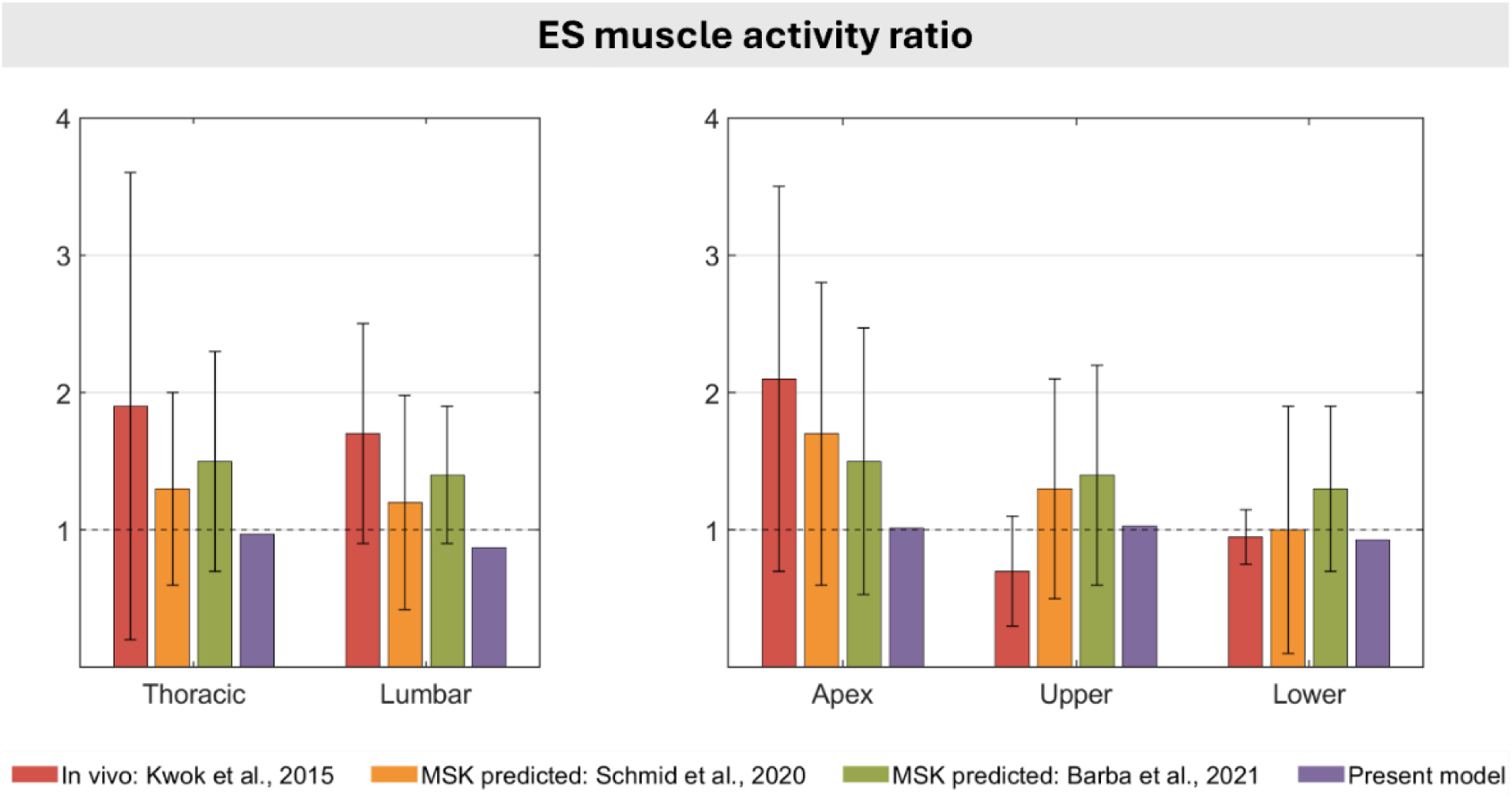
Convex-to-concave erector spinae (ES) activation ratios in the thoracic and lumbar regions (left), and at the curve apex and upper and lower ends of the curve (right), during upright standing.

**Fig. 4.**
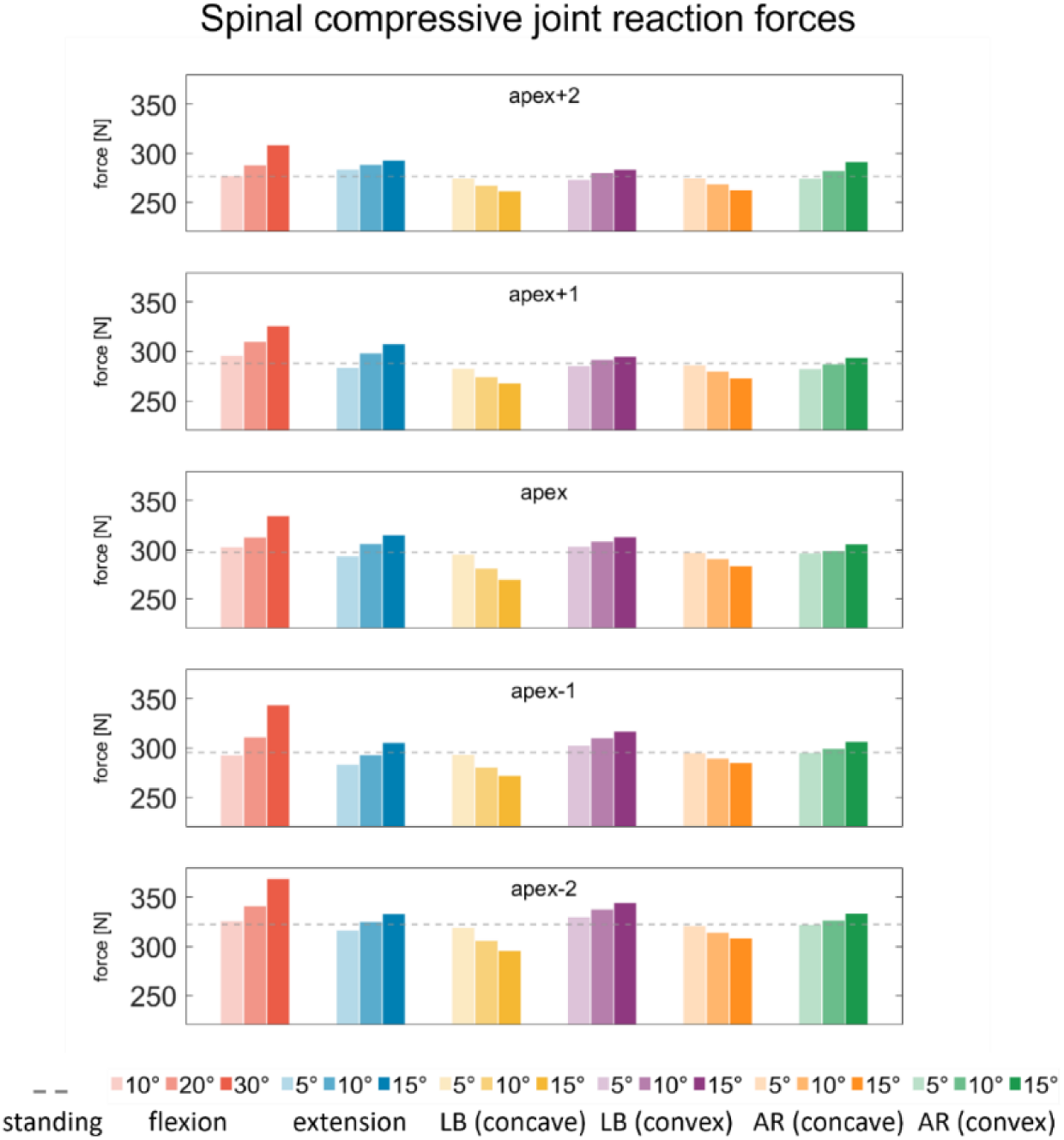
Spinal compressive joint reaction forces at the curve apex and two vertebral levels above and below the apex during simulated trunk postures. The dashed horizontal line indicates the reference condition of neutral upright standing.

**Fig. 5.**
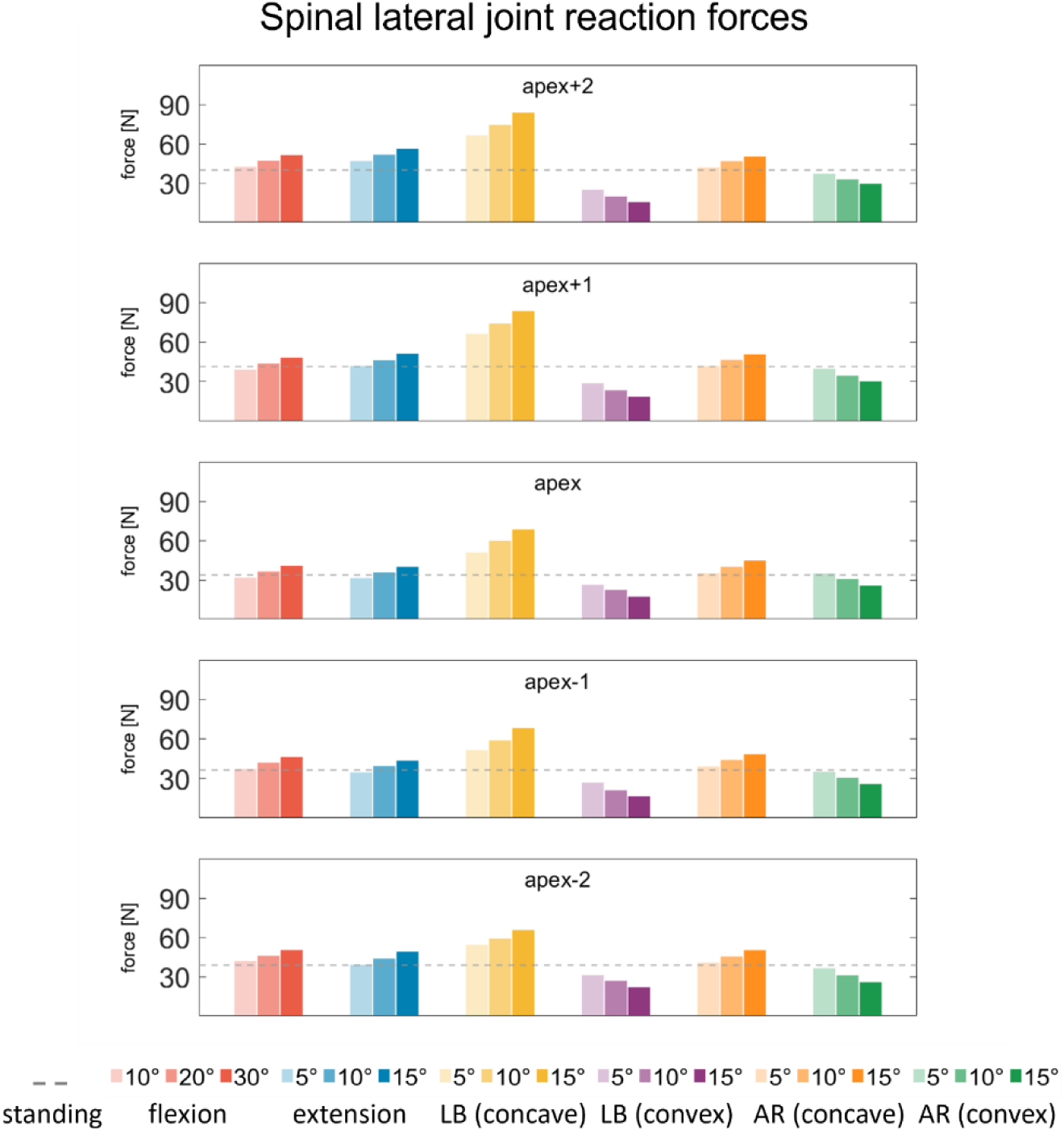
Spinal lateral joint reaction forces at the curve apex and two vertebral levels above and below the apex during simulated trunk postures. The dashed horizontal line indicates the reference condition of neutral upright standing.

### 3.2. Effect of trunk postures on muscle forces

The ES and MF muscles exhibited asymmetrical force patterns across the simulated trunk postures, with ES forces generally showing concave-side dominance and MF forces showing convex-side dominance (Fig. 6). Trunk flexion produced the highest muscle forces among all simulated postures, with both ES and MF forces increasing progressively with flexion angle. At 30° flexion, ES forces reached 200 N on the concave side and 180 N on the convex side, whereas MF forces reached 100 N on the convex side and 74 N on the concave side. During extension, both muscle groups maintained similar patterns of asymmetry, although the increase in muscle forces with increasing extension angle was comparatively moderate. During lateral bending, muscle forces decreased progressively with increasing concave bending and increased with increasing convex bending. At 15° concave bending, ES forces decreased to 76 N and 70 N on the concave and convex sides, respectively, while MF forces decreased to 53 N and 37 N. In contrast, at 15° convex bending, ES forces increased to 118 N and 88 N, while MF forces increased to 88 N and 57 N on the concave and convex sides, respectively. Axial rotation exhibited a similar asymmetrical muscle force pattern to lateral bending, although the overall magnitude of muscle forces was lower.

**Fig. 6.**
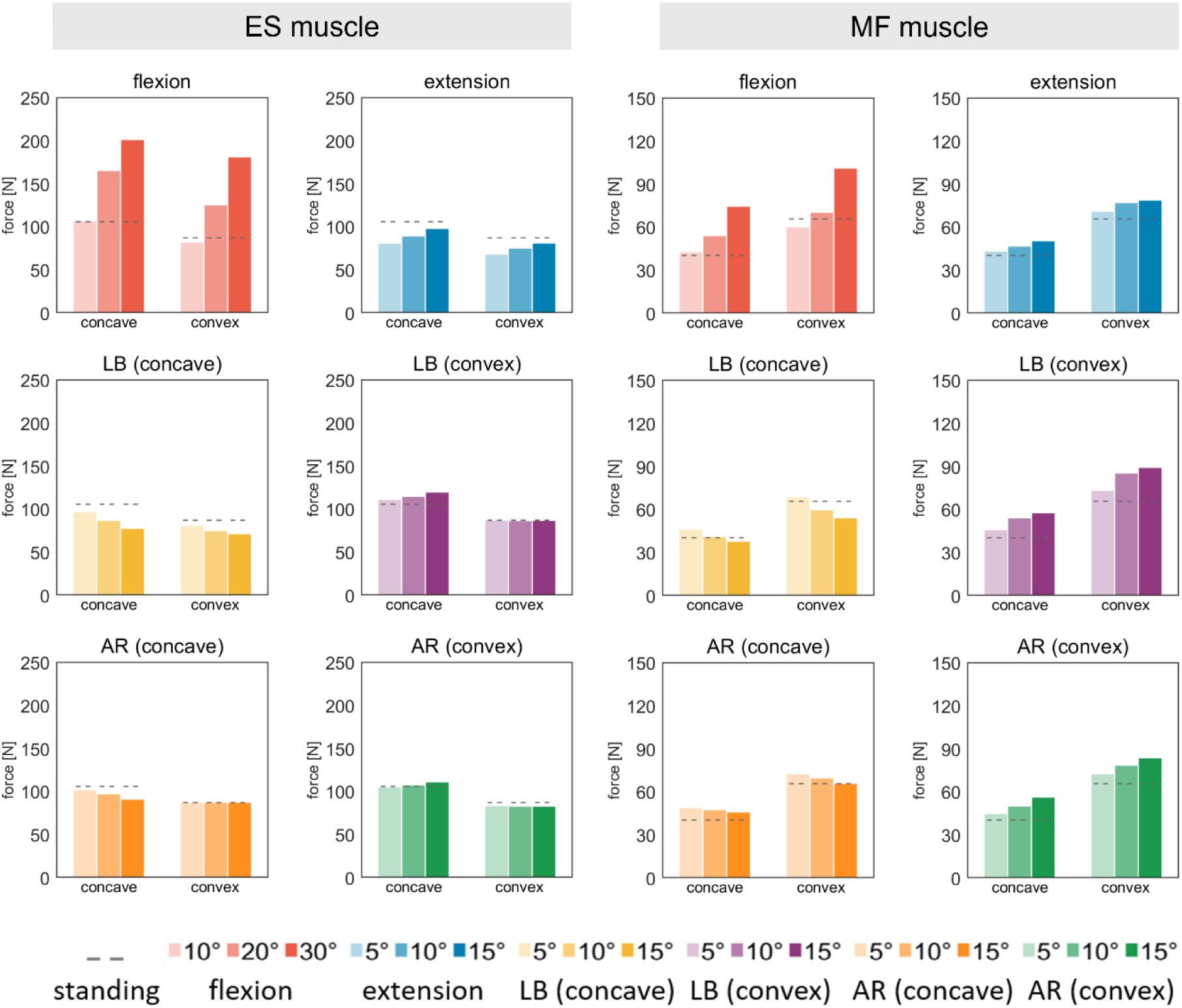
Muscle forces generated by the erector spinae (ES) and multifidus (MF) on the concave and convex sides of the curve during simulated trunk postures. The dashed horizontal line indicates the reference condition of neutral upright standing.

The level-wise distribution of muscle force asymmetry revealed distinct spatial patterns for the ES and MF muscles (Fig. 7). ES forces showed predominantly concave-side dominance, which was mostly pronounced below the curve apex and increased toward the lower lumbar levels. In contrast, MF forces showed predominantly convex-side dominance, particularly above the apex and toward the upper end of the curve. The overall spatial pattern of asymmetry remained largely consistent across activities within each muscle group. Among the simulated postures, flexion produced greater muscle force asymmetry than extension, while convex-side bending and rotation generally produced greater asymmetry than the corresponding concave-side postures.

**Fig. 7.**
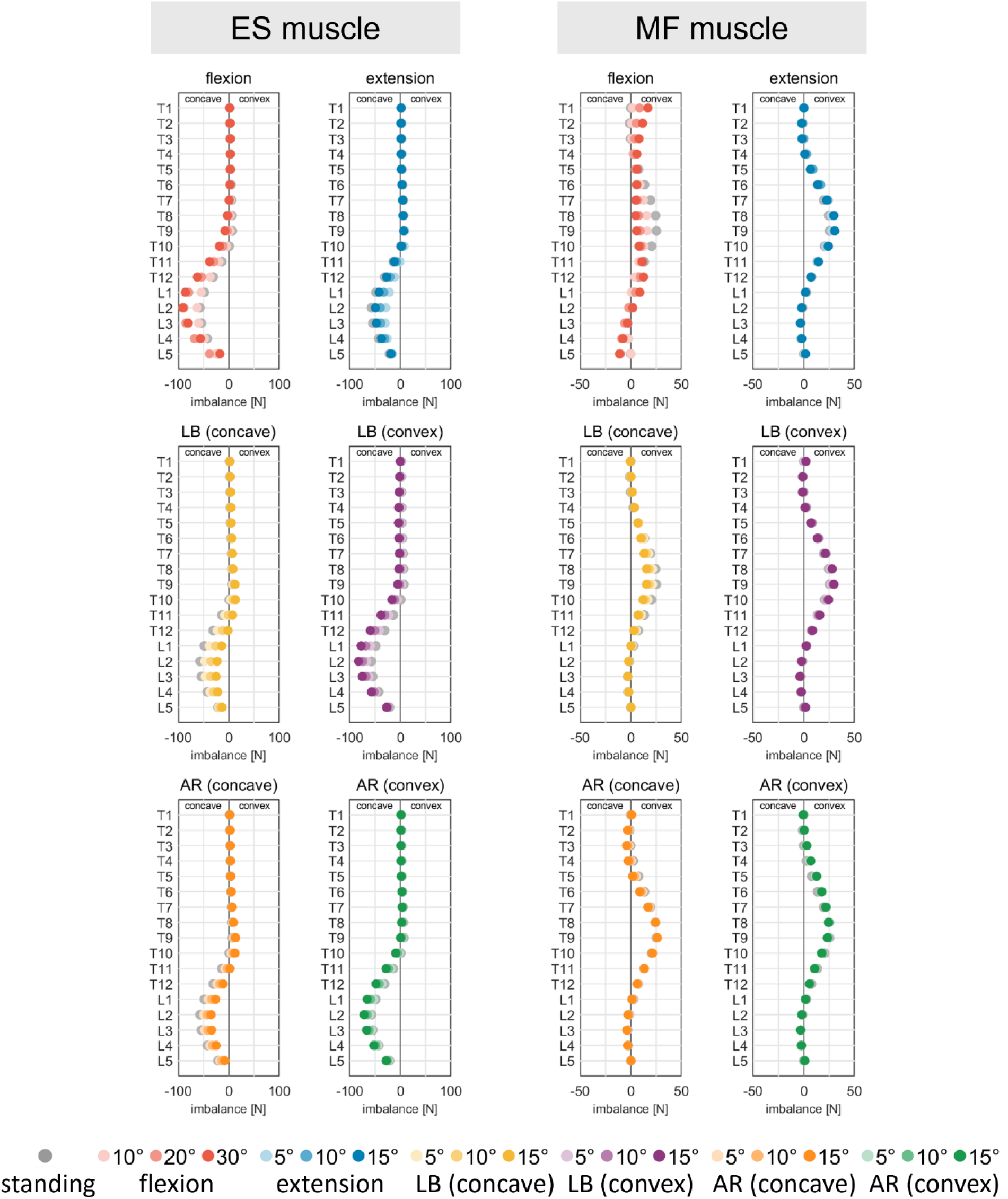
Muscle force imbalance (convex − concave) in the erector spinae (ES) and multifidus (MF) across simulated trunk postures at individual spinal levels. Negative values indicate concave-side dominance, whereas positive values indicate convex-side dominance.

### 3.3. Effect of trunk postures on intervertebral reaction forces

Trunk flexion consistently produced the highest compressive intervertebral forces among the simulated trunk postures (Fig. 4). Compared with neutral upright standing, compressive forces increased progressively with increasing flexion angle. At 30° flexion, compressive forces reached 337 N at the apex, 343 N at apex–1, and 372 N at apex–2, compared with 299 N, 296 N, and 324 N during upright standing, respectively. Extension produced comparatively smaller increases, reaching up to 312 N at the apex, 298 N at apex–1, and 303 N at apex+1 at 15° extension. Lateral bending exhibited a clear direction-dependent response. At 15° concave bending, compressive forces decreased to 280 N at the apex, 285 N at apex–1, and 290 N at apex–2, whereas 15° convex bending increased the corresponding forces to 320, 329, and 323 N, respectively. Axial rotation showed similar but less pronounced direction-dependent changes in compressive forces.

Compared with neutral upright standing, flexion and extension produced relatively small increases in lateral forces across the evaluated spinal levels (Fig. 5). In contrast, lateral bending produced substantially greater direction-dependent changes. At 15° concave bending, lateral forces increased to 87 N at apex+2, 84 N at apex+1, and 77 N at the apex, compared with 39 N, 41 N, and 32 N during upright standing, respectively. In contrast, 15° convex bending reduced the lateral forces to 22 N, 20 N, and 23 N, compared with 32 N, 34 N, and 39 N during upright standing, respectively. Axial rotation produced similar but less pronounced direction-dependent changes in lateral forces.

## 4. Discussion

This study investigated the influence of trunk posture on intervertebral loading and paraspinal muscle forces in an individual with AIS using a subject-specific MSK model. The patient-specific spinal deformity was incorporated into an age- and sex-matched adolescent healthy MSK model (Schmid, Burkhart, Allaire, Grindle, & Anderson, 2020), allowing the biomechanical response of the scoliotic spine to be evaluated across the simulated trunk postures (i.e., flexion, extension, lateral bending, and axial rotation). The findings demonstrated distinct posture-, direction-, and magnitude-dependent changes in intervertebral loading and paraspinal muscle forces, highlighting the importance of considering functional trunk postures when evaluating the biomechanical environment of the scoliotic spine.

The predicted ES muscle activation patterns were generally consistent with previous MSK modelling studies (Bassani et al., 2024; Schmid, Burkhart, Allaire, Grindle, Bassani, et al., 2020), although the present model predicted more symmetrical activation than the in vivo EMG measurements reported by (Kwok et al., 2015). The discrepancy between the predicted ES muscle activation and in vivo measurements may be attributed to inter-individual variations in curve type and severity among patient cohorts (Park et al., 2021). Additionally, the present model did not incorporate the muscle force–length–velocity relationship during muscle activation prediction due to the limited understanding of how AIS affects muscle physiology. This modelling simplification may have also contributed to the observed differences between predicted and EMG-based muscle activations. Nonetheless, the predicted activation ratios were within the physiological range reported in previous in vivo (Kwok et al., 2015) and MSK modelling studies (Bassani et al., 2024; Schmid, Burkhart, Allaire, Grindle, Bassani, et al., 2020).

The present study demonstrated consistent asymmetrical force patterns in the ES and MF muscles across all simulated trunk postures, with concave-side dominance in the ES and convex-side dominance in the MF. The concave-side dominance of the ES muscle may reflect its primary role as a global extensor and stabilizer (Hofste et al., 2020; Shima et al., 2022), generating greater forces on the concave side to counteract the tendency of the spine to collapse toward the convex side under gravitational loading. In contrast, the convex-side dominance of the MF muscle may reflect its role as a local stabilizer (MacDonald et al., 2006; Oshikawa et al., 2021), providing segmental stabilization, particularly in the vicinity of the curve apex, where precise control is required to maintain intervertebral alignment and vertebral stability. Together, these findings indicate that the altered spinal alignment in AIS leads to side-dependent muscle recruitment to maintain mechanical stability.

The consistent asymmetrical muscle force patterns observed in the present study are in agreement with previous experimental (Farahpour et al., 2014, 2015; Gou et al., 2024; Kwok et al., 2015; W. Wang et al., 2022) and MSK modelling studies (Barba et al., 2021; Bassani et al., 2024; Rauber et al., 2024; Schmid, Burkhart, Allaire, Grindle, Bassani, et al., 2020) in AIS, which have reported similar patterns of asymmetric paraspinal muscle activation. However, direct comparison with previous findings remains limited by differences in curve characteristics, including curve type, severity, and apex location, as well as differences in modelling approaches and patient characteristics.

The present simulations demonstrated that intervertebral reaction forces in AIS were strongly dependent on trunk posture. Trunk flexion produced the greatest compressive loading, with forces increasing progressively with flexion angle. During 30° flexion, compressive forces increased relative to upright standing at the apex (337 vs. 299 N), apex−1 (343 vs. 296 N), and apex−2 (372 vs. 324 N). This increase is likely attributable to the anterior displacement of the upper body mass, which increases the external flexion moment and the muscular demand required to stabilize the trunk. In contrast, trunk extension reduced compressive loading, likely due to a smaller external bending moment and reduced muscular stabilization demand.The present simulations demonstrated that intervertebral reaction forces in AIS are highly sensitive to both the type and magnitude of trunk posture. Among the simulated postures, trunk flexion resulted in the greatest mechanical demand, primarily due to the anterior displacement of the upper body mass relative to the spinal column, which increases the flexion moment arm and requires greater muscular stabilization. Consequently, compressive forces increased progressively with flexion angle, reaching 337 N at the apex, 343 N at apex−1, and 372 N at apex−2 during 30° flexion, compared with 299 N, 296 N, and 324 N during neutral upright standing, respectively. Similarly, concave(right)-side bending increased lateral loading relative to upright standing, particularly at and above the curve apex (apex+2: 87 vs. 39 N; apex+1: 84 vs. 41 N; apex: 77 vs. 32 N at 15° concave bending and upright standing, respectively). In contrast, trunk extension resulted in lower compressive loading, likely because the gravitational force vector becomes more closely aligned with the spinal axis, reducing external bending moments and the associated muscular stabilization demand.

Lateral bending produced clear direction-dependent changes in intervertebral loading, reflecting the asymmetric biomechanical response of the scoliotic spine. During 15° concave-side bending, lateral forces increased relative to upright standing at apex+2 (87 vs. 39 N), apex+1 (84 vs. 41 N), and the apex (77 vs. 32 N), likely due to greater geometric asymmetry and uneven load transmission across the curve. In contrast, convex-side bending reduced lateral forces, possibly by partially straightening the scoliotic curve and promoting more vertical load transmission. These trends are consistent with (Bassani et al., 2024), who similarly reported increased lateral forces during concave-side bending and decreased forces during convex-side bending, particularly at the upper curve levels.

Several limitations of the present study should be acknowledged. First, the present model did not include passive structures, such as ligaments and joint capsules, due to the lack of AIS-specific stiffness properties (Barba et al., 2021). Second, the model did not incorporate subject-specific muscle architecture. The use of generic muscle properties may reduce the accuracy of muscle force predictions, particularly in AIS, where spinal deformity can affect muscle orientation, cross-sectional area, and moment arms (Becker et al., 2023; Duncombe et al., 2025). Incorporating subject-specific anatomical data derived from imaging modalities such as MRI may improve the predictive capability of future simulations (Anderson et al., 2012; Keenan et al., 2014). Moreover, muscle forces were estimated using a static optimization approach, which assumes an optimal distribution of muscle forces and may not adequately represent altered or compensatory neuromuscular control strategies associated with AIS (Rauber et al., 2024). Finally, the intervertebral motion distributions prescribed for the simulated trunk postures were derived from healthy populations, as AIS-specific intervertebral kinematic constraints are currently unavailable.

## 5. Conclusions

This study investigated the influence of trunk posture on intervertebral loading and paraspinal muscle forces in an individual with AIS using a subject-specific MSK model. Trunk flexion produced the greatest compressive loading, whereas lateral bending generated pronounced direction-dependent changes in lateral forces, with concave-side bending increasing and convex-side bending reducing lateral loading. Paraspinal muscle forces were consistently asymmetric, with predominantly concave-side ES and convex-side MF dominance. Overall, these findings highlight the combined influence of spinal deformity and trunk posture on segmental spinal loading and muscle force generation in AIS. However, the clinical significance of these altered loading patterns, particularly their potential relationship with curve progression, joint degeneration, and back pain, remains unclear. Future studies should investigate how alterations in spinal loading and paraspinal muscle activity relate to curve progression and clinical outcomes, thereby improving the understanding of the biomechanical mechanisms underlying AIS.

## CRediT authorship contribution statement

**Rounak Bhattacharya:** Data Curation, Formal Analysis, Investigation, Methodology, Software, Validation, Visualisation, Writing – original draft. **Bhavuk Garg:** Conceptualisation, Methodology, Patient Recruitment Supervision, Writing – review & editing. **Rajesh Malhotra:** Conceptualisation, Methodology, Patient Recruitment Supervision, Writing – review & editing. **Rajdeep Ghosh:** Methodology, Investigation, Writing – review & editing. **Anoop Chawla:** Conceptualisation, Methodology, Resources, Supervision, Writing – review & editing. **Kaushik Mukherjee:** Conceptualisation, Methodology, Resources, Supervision, Writing – review & editing.

## Acknowledgement and Funding

The authors are grateful to the Indian Institute of Technology Delhi and the All India Institute of Medical Sciences, New Delhi, for providing the facilities and infrastructure required to conduct this study.

## Statements of ethical approval

Ethical approval was obtained from the Institute Ethics Committee (IEC) of the All India Institute of Medical Sciences, New Delhi, India, under approval number IEC-248/04.03.2022.

